# Associations between prenatal distress, mitochondrial health, and gestational age: findings from two pregnancy studies in the USA and Turkey

**DOI:** 10.1101/2024.10.16.618719

**Authors:** Qiuhan Huang, David Shire, Fiona Hollis, Sameera Abuaish, Martin Picard, Catherine Monk, Elif Aysimi Duman, Caroline Trumpff

**Affiliations:** Division of Behavioral Medicine, Department of Psychiatry, Columbia University Irving Medical Center, New York, NY, USA; Department of Pharmacology, Physiology and Neuroscience, University of South Carolina School of Medicine, Columbia, SC, USA; Department of Basic Sciences, College of Medicine, Princess Nourah Bint Abdulrahman University, Riyadh, Kingdom of Saudi Arabia; Department of Neurology, H. Houston Merritt Center, Neuromuscular Medicine Division, Columbia University Irving Medical Center, New York, NY, USA; New York State Psychiatric Institute, New York, NY, USA; Robert N Butler Columbia Aging Center, Columbia University Mailman School of Public Health, New York, NY, USA; Department of Obstetrics and Gynecology, Columbia University Irving Medical Center, New York, NY, USA; Department of Molecular Biology and Genetics, Faculty of Engineering and Natural Sciences, Acibadem University, Istanbul, Turkey; Institute of Natural and Applied Sciences, Acibadem University, Istanbul, Turkey

## Abstract

**Objective:** This study examined associations between mitochondrial markers—circulating cell-free mitochondrial DNA (cf-mtDNA) and Growth Differentiation Factor-15 (GDF15)—with maternal distress and pregnancy outcomes.

**Method:** Participants were drawn from two pregnancy studies, EPI (N=187, USA) and BABIP (N=198, Turkey). Plasma cf-mtDNA and GDF15 levels were quantified using qPCR and ELISA assays.

**Results:** Plasma cf-mtDNA levels did not significantly vary across pregnancy, while plasma GDF15 levels increased from early to late pregnancy and decreased postpartum. Late 2nd trimester plasma GDF15 was negatively correlated with pre-pregnancy BMI (p=0.035) and gestational age (p=0.0048) at birth. Early 2nd trimester maternal distress was associated with lower cf-mtDNA (p<0.05) and a trend for lower GDF15. Higher pre-pregnancy BMI and late-pregnancy maternal distress were linked to smaller postpartum GDF15 declines in EPI (p<0.05).

**Conclusion:** This study reveals distinct plasma cf-mtDNA and GDF15 patterns during the perinatal period, linking mitochondrial markers to maternal distress and pregnancy outcomes.

## Introduction

Perinatal maternal psychological distress, such as perceived stress, anxiety, and depression, has long been associated with adverse pregnancy outcomes including increased risk of preeclampsia (1, 2), spontaneous abortion, (3, 4) and shorter gestational age (5, 6). However, the biological mechanisms underlying these effects are still largely unknown. Mitochondria produce energy essential for stress adaption (7) and mitochondrial biology represents a potential intersection point between psychosocial experiences and their biological embedding ((8–15) for a review see (16)). Pregnancy is associated with a progressive increase in energy expenditure ((17–19), for a review see (20)). The psychological stress response recruits energy-demanding cellular and physiological processes that can compete with growth-related processes, causing energy constraints that may contribute to the biological embedding of stress and adversity across the lifespan (21). Thus, the energetically demanding period of pregnancy may compound pre-existing vulnerability, making it a time where mothers are particularly sensitive to the detrimental effects of psychological and metabolic stress on mitochondrial biology, potentially influencing pregnancy outcomes.

The mitochondrion is the only mammalian organelle besides the nucleus to contain its own genome. Each mitochondrion contains multiple copies of the 16.6kb-long circular mitochondrial DNA (mtDNA) (22, 23), which is consistently detectable outside of cells in most bodily fluids, including blood, as cell-free mitochondrial DNA (cf-mtDNA) (24). Under conditions of energetic stress, cf-mtDNA can be released into circulation, thereby acting as a biomarker for mitochondrial stress and signaling. cf-mtDNA levels in blood are elevated in several disease conditions, such as sepsis (25–27), cancer (for a review see (28)), infections (for a review see (29)), autoimmune disease (30–32), and psychopathology ((33, 34), for a review, see: (35)). In healthy non-pregnant individuals, levels of circulating cf-mtDNA are elevated following acute psychological stress (36, 37). While nothing is known about the connection between psychosocial stress, prenatal maternal distress, and cf-mtDNA in pregnancy, abnormal levels of cf-mtDNA have been found in pregnant women with preeclampsia (38) and gestational diabetes (39), suggesting a connection between adverse pregnancy outcomes and energetic stress.

Growth differentiation factor 15 (GDF15) is another emerging marker of energetic stress implicated in pregnancy. GDF15 is a member of the TGFβ super family that is released to modulate energy metabolism in response to mitochondrial and metabolic stress (40, 41). Throughout the human body, GDF15 is most highly expressed in placenta tissues (GTEx consortium (42)). The sole known receptor of GDF15, GFRAL, is located in the hindbrain, where the GDF15-GFRAL complex regulates whole-body energy homeostasis (41, 43) and supports energy mobilization (44). In pregnancy, circulating levels of serum GDF15 can increase up to 200-fold during the 3^rd^ trimester compared to non-pregnant postpartum state (45). Altered GDF15 levels in pregnancy have been associated with miscarriage (46), preeclampsia (47–50), gestational diabetes (50) and recently causally linked to hyperemesis gravidarum (51). Pregnancy triggers an unparalleled elevation in blood GDF15 signaling onto the brain to alter physiology (44, 52). No prior studies have investigated the interplay between GDF15 and prenatal maternal distress. Psychiatric disorders such as major depressive disorder have been associated with elevated levels of GDF15 in non-pregnant populations (53–55), and acute psychological stress exposure also increases circulating levels of GDF15 (56). These findings position GDF15 as an emerging marker of i) mitochondrial and energetic stress, ii) normal pregnancy physiology, iii) mental stress and psychopathology.

Taken together, cf-mtDNA and GDF15 are two emerging biomarkers that can offer insights into how energetic stress and maternal distress could converge to impact pregnancy outcomes. Here, we examined how levels of cf-mtDNA and GDF15 change across the perinatal period and their interplay with maternal perinatal characteristics, psychological distress (measured by perceived stress, anxiety and depression) and pregnancy outcomes in two pregnancy studies from the USA (EPI study) and Turkey (BABIP study).

## Results

We leveraged data and samples from two longitudinal pregnancy studies in the USA (EPI, N=187 (57)) and Turkey (BABIP, N = 198 (58)). Blood samples and psychological assessments were collected at four timepoints in EPI (early 2nd trimester to postpartum) and two timepoints in BABIP (late 2nd to 3rd trimester) (Figure 1A). Consequently, data from the early 2nd trimester and postpartum were only available in EPI. Due to recruitment challenges during early pregnancy, blood collection difficulties, and postpartum dropouts, no participants had measurements from all four time points. Given the non-normal distribution of cf-mtDNA and GDF15 levels, as well as the uneven sample sizes, non-parametric signed-rank Wilcoxon paired t-test was selected to maximize the use of available data. Demographic characteristics of the two studies are summarized in Table 1. On average, American EPI participants were significantly younger, less educated, and had a higher BMI compared to Turkish BABIP participants (ps<0.0001).

**Figure 1.**
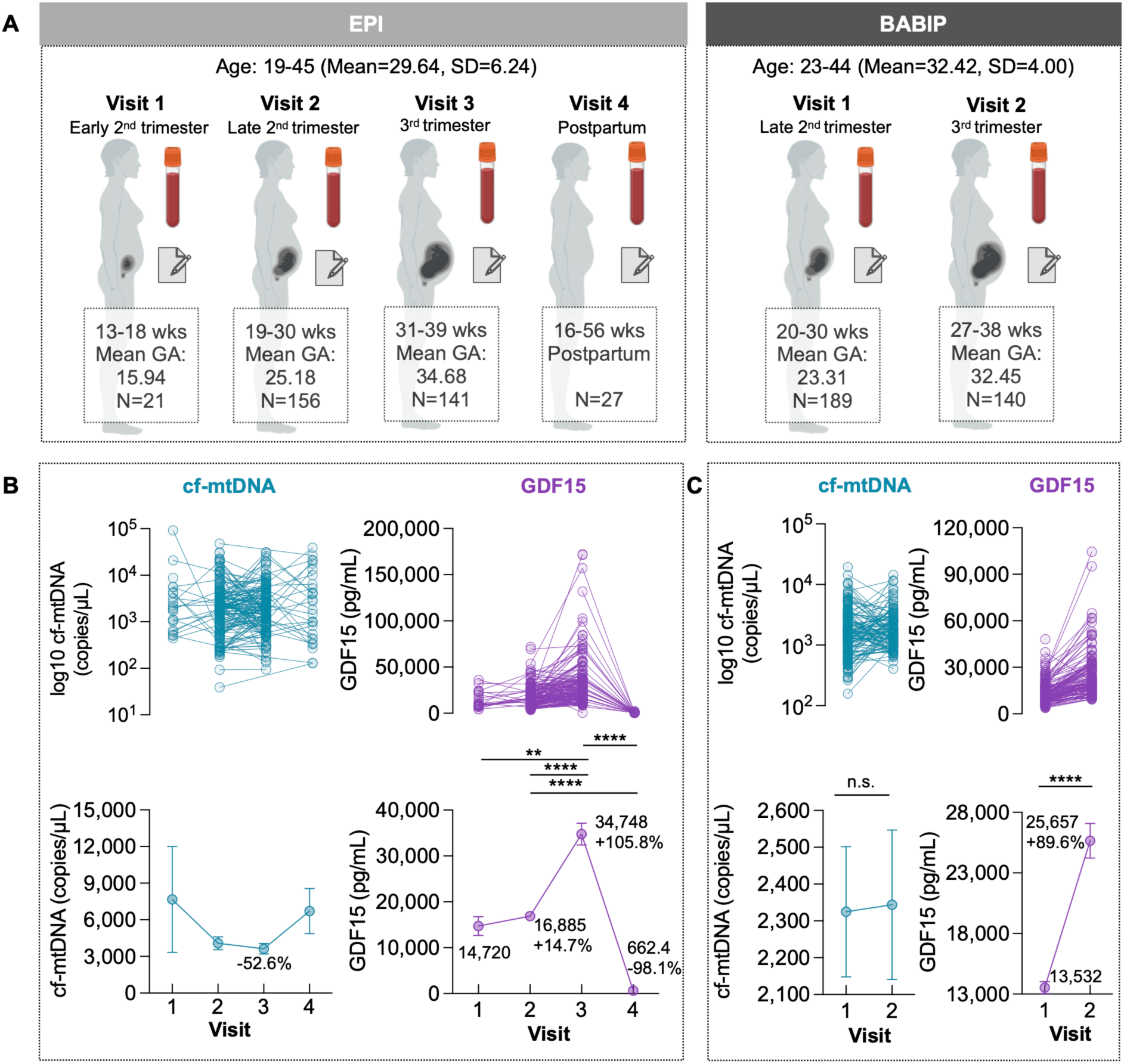
Maternal plasma cf-mtDNA and GDF15 levels across pregnancy. (**A**) Schematic of EPI (left) and BABIP (right) experimental design. cf-mtDNA and GDF15 concentrations were measured from plasma sampled across pregnancy during the early second trimester (13-18 weeks, EPI), late second trimester (19-28 weeks, EPI; 20-30 weeks BABIP) third trimester (32-39 weeks EPI; 27-38 weeks BABIP), and postpartum (16-56 weeks postpartum EPI). Participants completed questionnaires collecting information on prenatal distress at each visit. (**B**) Individual trajectories (top) and group average (bottom) of plasma cf-mtNDA and GDF15 levels in EPI, cf-mtDNA percent change from visit 1 to visit 3 is shown in the graph, GDF15 concentration at each visit and the percent change from the previous visit is shown in the graph. (**C**) Same as in (**B**) in BABIP. (B,C) Data shown as mean ± SEM. GA stands for gestational age. In EPI, no participants had data for all four time points. Given the non-normal distribution of cf-mtDNA and GDF15 levels and uneven sample sizes, pairwise comparisons were used to maximize data use and provide clear insights while minimizing the impact of missing data and small samples. P-values from non-parametric signed-rank Wilcoxon paired t-test *p<0.05, **p<0.01, ***p<0.001, ****p<0.0001.

### cf-mtDNA and GDF15 trajectories across pregnancy

In contrast with a prior study (59), we did not find evidence of significant variation in plasma cf-mtDNA across pregnancy in either study (Figures 1B and 1C, left). GDF15 is most highly expressed in decidual stromal cells of placenta, (GTEx consortium ((42), see Figure S1) and it can be hypothesized that as the placenta grows, GDF15 levels would increase. In both studies, we found evidence of a continuous increase in plasma GDF15 levels from early to late pregnancy. In EPI, plasma GDF15 levels gradually increased from early to mid-pregnancy (+14.7%), doubled from mid to late pregnancy (+105.8%), and dropped sharply after postpartum (−98.1%) to levels comparable with non-pregnant healthy controls (Figures 1B, right). In BABIP, plasma GDF15 showed similar magnitude of increase from mid to late-pregnancy (+87.2%) (Figure 1C, right).

In EPI, cf-mtDNA levels measured in the late 2nd and 3rd trimesters showed a moderate positive correlation (Figure S2A), which was not observed in BABIP (Figure S2B). For both studies, we observed strong positive correlations in GDF15 levels measured in the late 2^nd^ and 3^rd^ trimesters (rs=0.49-0.65, p<0.0001), confirming the trait nature of this biomarker. No correlation was found between cf-mtDNA and GDF15 measured within the same visit or across different visits (Figures S2C-F).

### cf-mtDNA, GDF15 and maternal characteristics

Next, we investigated the association between cf-mtDNA, GDF15, and maternal characteristics. There were no associations between plasma cf-mtDNA, maternal age, and pre-pregnancy BMI in either study (Supplemental Table 1). In EPI, plasma GDF15—the most significantly upregulated protein in human aging (60) —was not associated with maternal age during pregnancy (Supplemental Table 2). The expected positive correlation between age and circulating GDF15 levels emerged in postpartum (16-56 weeks, Figure S3A, r=0.47, p=0.016). In BABIP, we found a modest negative correlation between 3^rd^ trimester plasma GDF15 levels and maternal age (Figure S3B, r=-0.18, p=0.049). Regarding pre-pregnancy BMI, higher values were associated with lower plasma GDF15 levels in EPI during the late 2^nd^ trimester (Figure S3C [left], r=-0.18, p=0.035) and 3^rd^ trimesters (Figure S3C [right], r=-0.15, p=0.070). In BABIP, a similar pattern was found in the late 2^nd^ trimester (Figure S3D [left], r=-0.13, p=0.088), but not in the 3^rd^ trimester (Figure S3D [right]).

### cf-mtDNA, GDF15, neonatal characteristics and perinatal complications

In addition to maternal characteristics, we explored whether cf-mtDNA and GDF15 levels differ according to neonatal sex, gestational age at birth, or adverse pregnancy outcomes such as preeclampsia, preterm birth, and gestational diabetes.

In both studies, we found no significant differences in maternal plasma cf-mtDNA levels based on neonatal sex (Supplemental Table 3). Similarly, plasma cf-mtDNA levels at any sampling point were not significantly associated with gestational age at birth. (Figures 2A-B). In EPI, participants who developed preeclampsia (n=6) tended to present lower levels of plasma cf-mtDNA in the late 2^nd^ and 3^rd^ trimesters compared to those without, but the difference did not reach statistical significance (Supplemental Table 4). The sample size of participants with preeclampsia (n=2) was insufficient to investigate this question in BABIP. Preterm birth (<37 weeks) was associated with elevated plasma cf-mtDNA levels in the late 2nd trimester in EPI participants (n=10, p=0.044), but no such difference was observed in BABIP (n=8, Supplemental Table 4). Consistent with a previous study that reported elevated circulating cf-mtDNA levels in women with gestational diabetes mellitus (39), we found higher plasma cf-mtDNA levels in the late 2^nd^ trimester (p=0.022) in EPI and in the 3^rd^ trimester (p=0.013) in BABIP (Supplemental Table 4).

**Figure 2.**
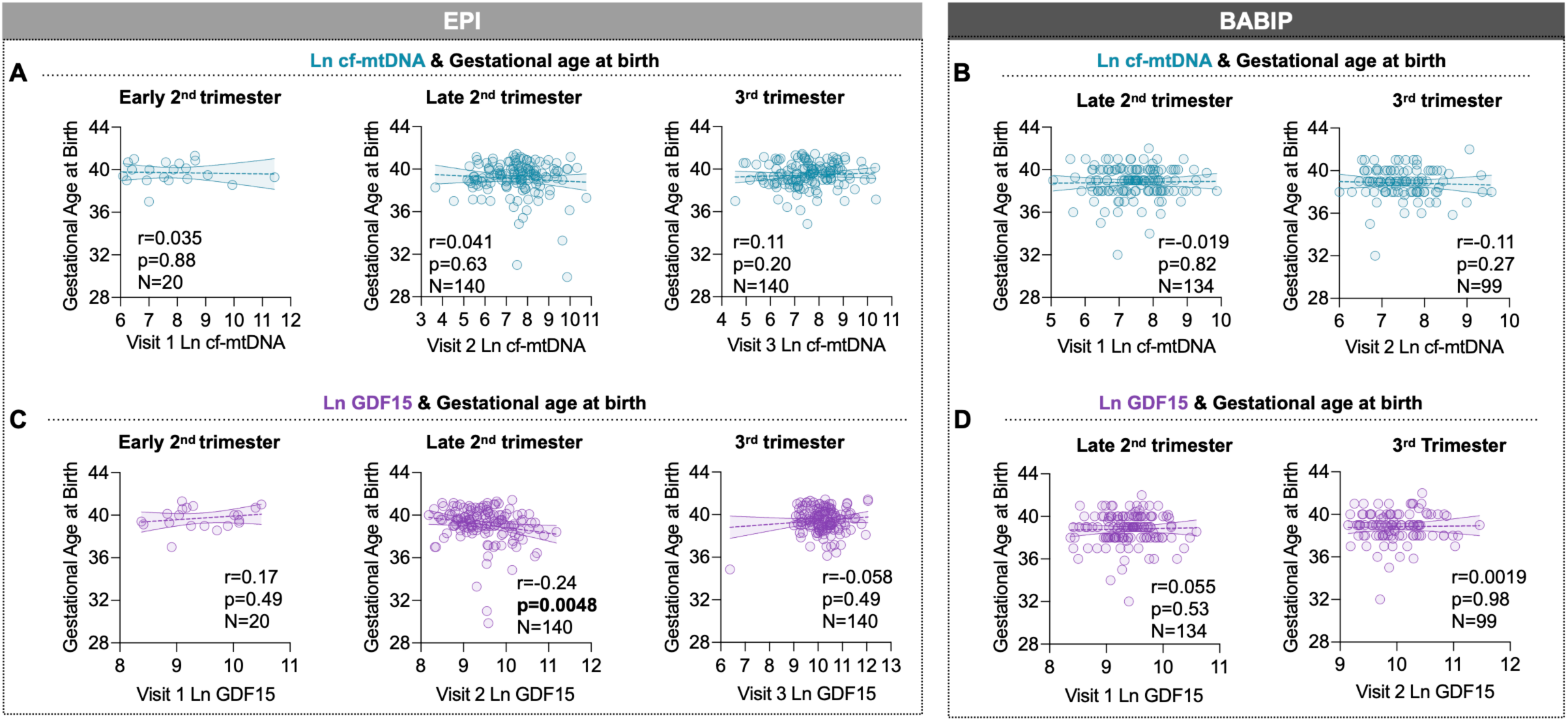
Association between maternal plasma cf-mtDNA and GDF15 levels and gestational age at birth. Associations between gestational age at birth and cf-mtDNA at each visit in EPI (**A**) and BABIP (**B**). Associations between gestational age at birth and GDF15 at each visit in (**C**) EPI and (**D**) BABIP. 13 EPI participants and 2 BABIP participants were removed from the analysis and graphs since they completed their late 2nd trimester visit in their 3rd trimester. See Supplemental Figure S4 for cf-mtDNA and GDF15 data shown in raw values. P-values and effect sizes from Spearman’s rank correlation. *p<0.05, **p<0.01, ***p<0.001, ****p<0.0001.

Regarding GDF15, a previous study reported significant elevation in 1^st^ trimester serum GDF15 levels in pregnant women carrying female offspring (45). Similarly, in the late 2^nd^ and 3^rd^ trimesters of both EPI and BABIP with more robust sample sizes, we observed higher average GDF15 levels in the 2^nd^ and 3^rd^ trimesters in women carrying female offspring compared to those carrying male offspring, although these differences did not reach statistical significance (Supplemental Table 3). A negative correlation was found between gestational age at birth and late 2^nd^ trimester GDF15 levels in EPI (r=-0.24, p=0.0048) but not in BABIP (r=0.055, p=0.53) (Figures 2C-D). In EPI, we observed that participants who developed preeclampsia (n=6) showed lower levels of plasma GDF15 in late 2^nd^ trimester (p=0.017, Supplemental Table 4). The sample size of participants with preeclampsia (n=2) was insufficient to investigate this question in BABIP. Unlike cf-mtDNA, GDF15 did not show any significant difference by maternal gestational diabetes status in EPI or BABIP (Supplemental Table 4). Further, maternal plasma GDF15 levels did not differ between preterm and full-term pregnancies in EPI (n=10) or BABIP (n=8) (Supplemental Table 4), although results should be considered with caution given that the number of neonates in the preterm group was too low.

### cf-mtDNA and GDF15 and maternal prenatal distress

In both studies, no significant associations were found between maternal plasma cf-mtDNA levels and perceived stress, anxiety or depressive symptoms in late 2^nd^ and 3^rd^ trimesters (Supplemental Table 1, for cf-mtDNA levels based on clinical cut-offs see Supplemental Table 5). Early 2^nd^ trimester data, available only in EPI, revealed that higher depressive symptoms (Figure 3A [top], r=-0.56, p=0.032) and higher perceived stress (Figure 3B [top], r =-0.72, p=0.0031) were associated with lower plasma cf-mtDNA levels. In the early 2^nd^ trimester, there was also a trend for a negative association between anxiety symptoms and plasma cf-mtDNA levels (Figure 3C [top], r=-0.51, p=0.055).

**Figure 3.**
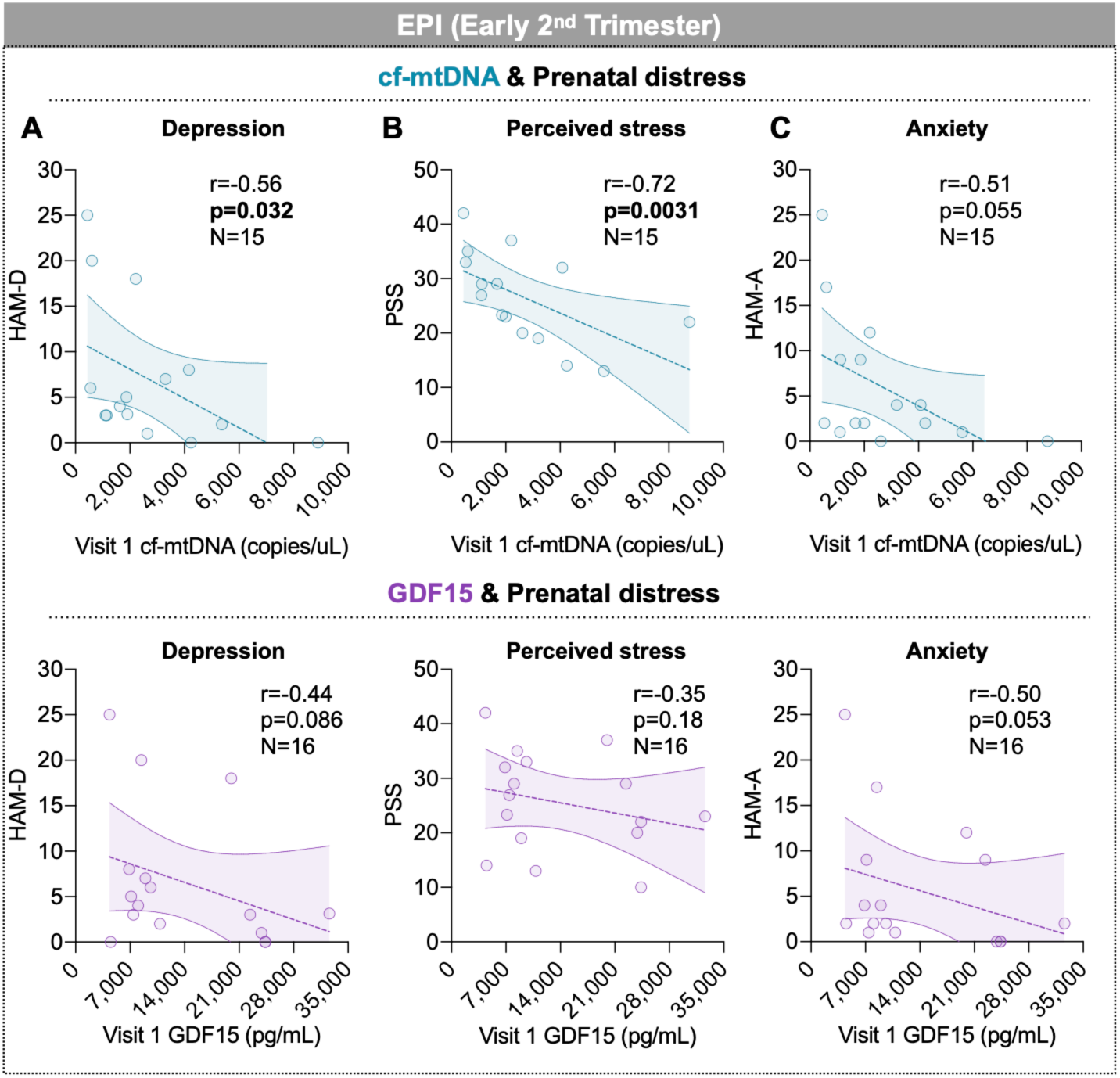
Associations between maternal plasma cf-mtDNA and GDF15 levels and maternal psychological distress in early pregnancy (available only in EPI study). Association between plasma cf-mtDNA (top) and GDF15 (bottom) levels in the early 2nd trimester (13-18 weeks) and (**A**) depressive symptoms assessed by the Hamilton depression rating scale (HAM-D), (**B**) psychological stress assessed by perceived stress scale (PSS), and (**C**) anxiety symptoms assessed by the Hamilton anxiety rating scale (HAM-A) in EPI. P-values and effect sizes from Spearman’s rank correlation. *p<0.05, **p<0.01, ***p<0.001, ****p<0.0001

Similar to cf-mtDNA, in the early 2^nd^ trimester of the EPI study, we found negative trends between maternal GDF15 plasma levels and depressive symptoms (Figure 3A [bottom], r =-0.44, p=0.086), anxiety symptoms (Figure 3B [bottom], r=-0.50, p 0.052), and perceived stress (Figure 3C [bottom], r=-0.35, p=0.18). In both studies, no significant associations were found between maternal GDF15 levels and perceived stress, anxiety or depressive symptoms in late 2^nd^ and 3^rd^ trimesters (Supplemental Table 1, for GDF15 levels based on clinical cut off see Supplemental Table 5), a period where placenta-related release of GDF15 might dominate the signal.

### Change in GDF15 levels and maternal characteristics during pregnancy

Given the significant changes in GDF15 levels throughout the perinatal period, we investigated whether maternal characteristics and prenatal distress could account for the individual differences in GDF15 trajectories from pregnancy to postpartum, using data from the EPI study, which included postpartum time points (Supplemental Table 6). Interestingly, we found that higher pre-pregnancy BMI was associated with lower decline in GDF15 from 3^rd^ trimester to post-partum (Figure 4A, r=0.41, p=0.044), which can be interpreted as an impaired return to pre-pregnancy baseline state. Similarly, higher perceived stress and depressive symptoms in the 3^rd^ trimester were associated with lower decline in GDF15 from the 3^rd^ trimester to post-partum (Figures 4B-C, r=0.43, p=0.042; r=0.59, p=0.004, respectively). Altogether, these findings suggest that elevated prenatal metabolic stress and maternal distress may interact with post-pregnancy physiological recovery.

**Figure 4.**
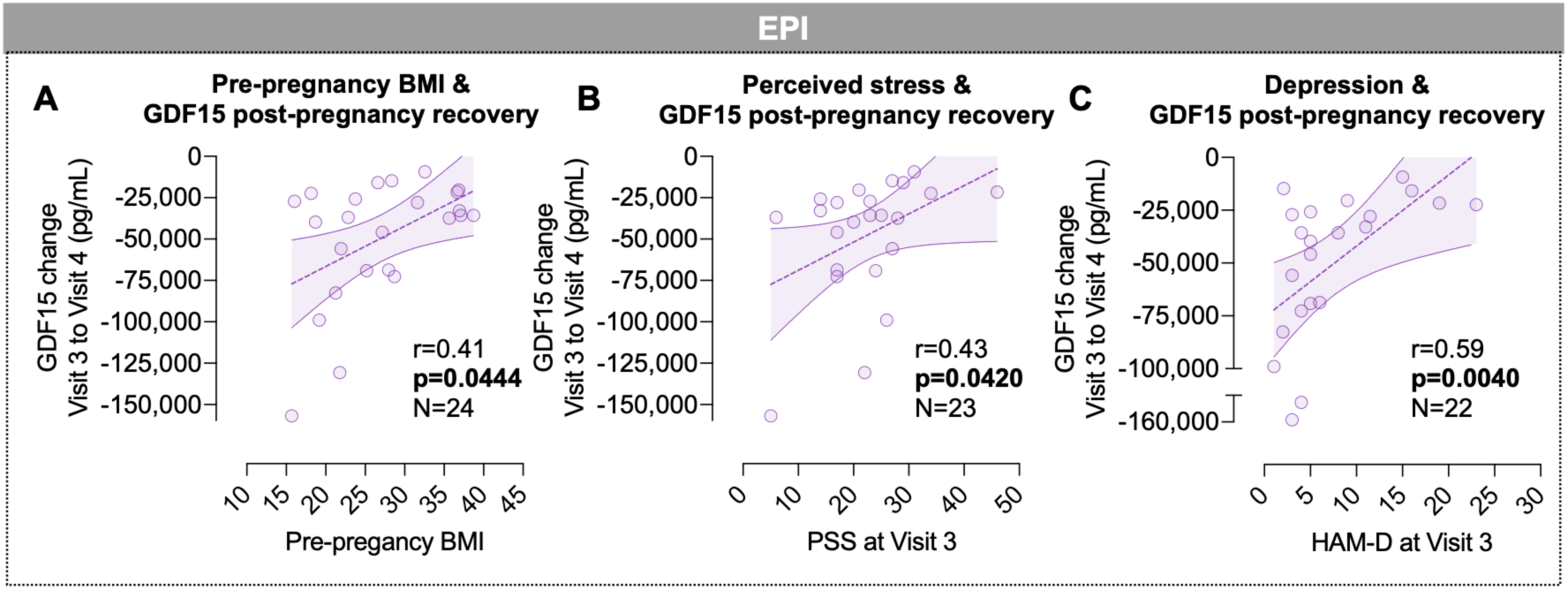
Associations between pre-pregnancy BMI, prenatal maternal distress and change in plasma GDF15 levels from late pregnancy to postpartum (available only in EPI study). Association between the change in plasma GDF15 from the 3r^d^ trimester (32-39 weeks) levels to postpartum (16-56 weeks after birth) levels and (**A**) pre-pregnancy BMI, (**B**) psychological stress assessed by perceived stress scale (PSS) during 3^rd^ trimester, and (**C**) depressive symptoms assessed by the Hamilton depression rating scale (HAM-D) during 3^rd^ trimester in EPI. P-values and effect sizes from Spearman’s rank correlation. *p<0.05, **p<0.01, ***p<0.001, ****p<0.0001

## Discussion

Leveraging longitudinal studies from two countries (USA and Turkey), we investigated the trajectories of perinatal plasma cf-mtDNA and GDF15 levels and their associations with maternal characteristics, prenatal distress, and pregnancy outcomes. We report patterns of both cf-mtDNA and GDF15 across gestation periods and trimester-specific associations with maternal and neonatal characteristics, some of which were consistent across the two studies.

Considering the lack of consistent literature, we first examined how plasma cf-mtDNA and GDF15 changes across the pregnancy. The lack of change in plasma cf-mtDNA levels across pregnancy contrasts with a recent longitudinal study of healthy pregnancies (n=32) that found a 1.7-fold increase in serum cf-mtDNA from early pregnancy (5-8 weeks) to late pregnancy (33-36 weeks)(59). This difference might stem from the different sampling intervals (i.e. having no early first trimester data) used in that study or the use of serum samples versus plasma (24). Regarding GDF15, we observed a significant increase in plasma levels from early to late pregnancy in both studies, followed by an average 98.1% decrease (range across women: 93.4-99.8%) postpartum relative to 3^rd^ trimester levels. Our findings are consistent with previous findings showing up to 100-fold increase in GDF15 levels comparing pregnant and non-pregnant populations (61, 62), and with another study reporting a 200-fold increase in GDF15 levels in the 3^rd^ trimester compared to post-partum (45). The dramatic decrease from 3^rd^ trimester to postpartum in our results also aligns with recent findings that show late pregnancy GDF15 levels were 172 times higher than those measured early postpartum(63). Our study therefore adds to a robust body of literature demonstrating that GDF15 is not only a marker of aging (60, 64) and mitochondrial and energetic stress (40, 41), but is also strongly associated with pregnancy course, rapidly returning to baseline after delivery.

In both studies, women with higher plasma GDF15 levels in the late 2^nd^ trimester also tended to have elevated levels in the 3^rd^ trimester, indicating moderate within-person stability. Plasma cf-mtDNA levels, on the other hand, showed within-person stability for EPI, but not BABIP, participants. No correlation was observed between cf-mtDNA and GDF15 levels, either measured within the same trimester or across different trimesters, suggesting that these two biomarkers may be at least partially regulated independently or influenced by individual-specific factors that have yet to be discovered.

Although GDF15 is strongly associated with aging (60, 64), we found no link between maternal age and GDF15 levels during pregnancy. In contrast, the anticipated positive association was found during the postpartum period. The lack of GDF15’s association with age during pregnancy might come from its role in pregnancy-specific metabolic and physiological adaptations (65), such as pregnancy-induced insulin resistance (66) and placental invasion (46). Interestingly, recent work suggests that epigenetic aging is accelerated during pregnancy and reversed during post-partum (67). Future work is needed to understand the interplay between plasma GDF15 levels and epigenetic aging trajectories during pregnancy.

Our results also link GDF15 to fetal development. Our finding showing that plasma GDF15 levels in mid pregnancy were negatively correlated with gestational age at birth in EPI highlights a potentially significant role of GDF15 in prenatal development and pregnancy outcomes. Interestingly, a recent study in preterm infants also found an inverse relationship between gestational age at birth and serum GDF15 levels at birth (68). Taken together, elevated GDF15 levels in mid-pregnancy could indicate pregnancy-related energetic stress, potentially leading to earlier delivery.

In pregnant populations, previous studies found a negative correlation between GDF15 levels and BMI (69), and lower GDF15 increase in pregnancy in obese participants compared to normal weight participants (66, 70). In line with these findings, we found that pre-pregnancy BMI and plasma GDF15 levels were inversely correlated during pregnancy, particularly in mid-pregnancy. Since higher BMI is often associated with metabolic stress (71), the inverse relationship between BMI and GDF15 in pregnancy may result from increased metabolic stress impairing GDF15 upregulation in pregnancy necessary for energy mobilization and metabolic adaptations (72). Additionally, we also found that higher pre-pregnancy BMI is associated with a lower decrease in GDF15 levels from 3^rd^ trimester to 4-14 months postpartum. This is in line with previous findings showing that the magnitude of the reversal of epigenetic aging observed from pregnancy to post-partum is lower in pregnant women with higher BMI (67). Taken together, these findings suggest that metabolic stress might increase the physiological load of pregnancy and alter post-pregnancy recovery.

A growing body of literature suggests that psychological stress and mood affect mitochondria biology (12, 73), including cf-mtDNA (35) and GDF15 levels (56) in non-pregnant adults. In this study, we found that higher perceived stress, depression, and anxiety symptoms are associated with lower cf-mtDNA levels in the early 2^nd^ trimester of pregnancy. While some studies have found cf-mtDNA levels to be elevated in depression (34), our findings are in line with a study showing cf-mtDNA levels to be lower in patients with major depressive disorder (74). However, the time-specific effect of prenatal distress on cf-mtDNA should be interpreted cautiously due to the small sample size.

In parallel, we found that women with higher levels of depressive and anxiety symptoms during early 2^nd^ trimester tended to have lower levels of plasma GDF15. This contrasts with previous studies indicating that GDF15 levels are elevated in psychopathological conditions in non-pregnant older adults (53). Early 2^nd^ trimester is a critical period for hormonal and physiological changes during pregnancy that may be supported partly by a GDF15 increase (75, 76). Our results suggest that the presence of psychological distress might counteract this elevation early in pregnancy, resulting in the observed lower levels of these markers in distressed pregnant women. If normal placental development is associated with increasing GDF15 release (independent of stress), then pregnancy-rise of GDF15 may be interpreted as a marker of normal progress and healthy pregnancy. Our observation that perceived stress and depressive symptoms were linked to a blunted post-partum decrease in GDF15 suggests that prenatal distress may interfere with post-pregnancy physiological recovery, perhaps through mechanisms or sources of GDF15 other than the placenta. Taken together, these findings uncover a novel potential link between prenatal distress and biomarkers of mitochondrial health and call for future studies investigating this relationship in larger studies.

This study is not without limitations. As discussed in detail in the methods, we used the R&D ELISA kits that has been shown to underestimate GDF15 levels in individuals carrying the H202D variant of the GDF15 gene (77). This limitation of the kit could have affected the results comparing absolute GDF15 levels between participants, but not the findings related to within-person. Despite the repeated-measures design, EPI had a small sample size in the early 2^nd^ trimester and postpartum. A key strength of this study is its unique cross-cultural approach. Although both studies are not directly comparable in terms of design and timepoints, the data from populations in the United States and Turkey reveals similar biological patterns across cultural contexts. This consistency enhances the robustness and broad relevance of mitochondrial markers like cf-mtDNA and GDF15 in relation to prenatal distress and pregnancy outcomes. However, this cross-cultural approach also raises challenges since the two studies involved different populations, sampling times, and assessments for depression and anxiety symptoms, which may affect the comparability of these measures. Future research should aim for more standardized assessments across diverse populations to further validate such findings.

## Conclusion

Using studies from two different countries, we describe distinct patterns in circulating plasma cf-mtDNA and GDF15 during pregnancy and their associations with maternal distress and pregnancy outcomes. Plasma cf-mtDNA levels showed no significant variation in either study, while GDF15 levels increased from early to late pregnancy and were negatively correlated with pre-pregnancy BMI in the late 2^nd^ trimester. Higher perceived stress and depressive symptoms were linked to lower cf-mtDNA levels during early pregnancy, indicating a potential early impact of maternal distress on mitochondrial markers. Maternal distress also influenced GDF15 trajectories, suggesting an interaction with post-pregnancy recovery. While they remain to be confirmed by larger studies, our findings from studies in two culturally distinct countries underscore cf-mtDNA and GDF15 as potential biomarkers for mitochondrial and psychosocial stress during pregnancy that can shed light on the biological mechanisms connecting maternal distress to adverse pregnancy outcomes.

## Methods

### 1. Participants

#### 1.1. EPI (USA)

Healthy pregnant women (N=187, ages 20–45; Mage=29.64, SDage=6.24) were recruited as part of the “Prenatal stress: the epigenetic bases of maternal and perinatal effects” (EPI) study during the years 2011–2016 through the Department of Obstetrics and Gynecology at Columbia University Medical Center as described previously (57). Exclusion criteria were multiparity, medication use, and tobacco or recreational drug use. Participants provided written informed consent prior to participating in the study. Participants completed their first visit either in the early (13-18 weeks) or late 2^nd^ trimester (19-30 weeks), depending on the time of recruitment, with subsequent visits occurring during the 3^rd^ trimester (31-39 weeks) and postpartum (16-56 weeks). 13 participants completed their late 2^nd^ trimester visit with gestational age greater than 28 weeks and were excluded from trimester-related analyses. During the study visit, oral and written consents were obtained by trained graduate assistants. Afterwards, participants completed questionnaires and a blood sample was collected by the study phlebotomist (Figure 1A, left). All procedures were approved by the Institutional Review Board of the New York State Psychiatric Institute/Columbia University Medical Center and all methods were performed in accordance with relevant guidelines and regulations.

#### 1.2. BABIP (Turkey)

Healthy pregnant women (N=198, ages 23-44; Mage=32.42, SDage=4.00) were recruited during the years 2018-2022 through doctors’ offices, flyers and online advertisements from Istanbul, Turkey as part of the “Bogazici Mother Baby Relationship Project” (BABIP) birth cohort as described previously (Duman et al., 2020). Exclusion criteria were multiparity and severe pregnancy complications. During lab visits, participants provided oral and written consents, completed questionnaires, and had blood samples collected by nurses. All procedures were approved by the Institutional Review Board of Bogazici University, where the study was initiated. Participants completed the first visit during their late 2^nd^ trimester (20-30 weeks) and the second visit during their 3^rd^ trimester (27-38 weeks). 2 participants completed their first visit with gestational age greater than 28 weeks and were excluded from trimester-related analyses. Information about pregnancy outcomes, such as gestational age, neonatal sex and perinatal complications were collected at one month after birth via online questionnaires (Figure 1A, right).

### 2. Psychosocial assessment and blood collection

#### 2.1. EPI (USA)

Prenatal distress in participants was evaluated at each visit using the Hamilton Depression Rating Scale (HAM-D)(78), the Hamilton Anxiety Rating Scale (HAM-A)(79), and the Perceived Stress Scale (PSS)(80). Blood was collected at each visit by the study phlebotomist using EDTA coated tubes. Plasma was isolated immediately after collection by centrifugation and was stored at −80°C until further processing.

#### 2.2. BABIP (Turkey)

Prenatal distress was evaluated at two prenatal visits during the late 2^nd^ trimester and 3^rd^ trimester using the Beck’s Depression Inventory-II (BDI-II)(81), the Center for Epidemiological Studies Depression (CESD)(82), the State-Trait Anxiety Inventory-State (STAI-S)(83), and the Perceived Stress Scale (PSS)(80). At each visit, blood samples were collected by nurses using EDTA coated tubes. Plasma was isolated immediately after collection by centrifugation and aliquots were stored at −80°C until they were transferred on dry ice to Columbia University Irving Medical Center for analysis.

### 3. Maternal characteristics and pregnancy outcomes

#### 3.1. EPI (USA)

Detailed information was collected during labor to comprehensively document neonatal and maternal outcomes. Recorded parameters included neonatal sex, gestational age at birth, and perinatal complications such as preeclampsia, preterm birth, and gestational diabetes.

#### 3.2. BABIP (Turkey)

Participants provided detailed information about perinatal complications, such as preeclampsia, preterm birth, and gestational diabetes, during the two prenatal visits as well as in the 1-month postpartum assessment. Pregnancy outcomes, such as gestational age at birth and neonatal sex were also recorded in the postpartum assessment.

### 4. GDF15 assays

For both studies, plasma GDF15 levels were quantified using a high-sensitivity ELISA kit (R&D Systems, DGD150) following the manufacturer’s instructions. Plasma samples were diluted with assay diluent (1:64 ratio for pregnancy samples, 1:4 ratio for postpartum samples) to maximize the number of samples within the dynamic range of the assay. Absorbance was gauged at 450nm, and concentrations were computed utilizing the Four Parameter Logistic Curve (4PL) model. Samples were run in duplicates on separate plates and the concentration for each sample was computed from the average of the duplicates. Samples with C.V.s larger than 15% were re-run. Samples with concentration above the dynamic range of the assay were rerun with 1:256 dilution with assay diluent. Standard curve (5 samples per plate) and plasma reference samples (3 samples per plate) were run with each individual assay and the inter-assay C.V. was monitored. All standard curves and references were overlaid on top of each other to monitor failed runs. Data-preprocessing and quality control measures was done using the R Software (version 4.2.2).

### 5. cf-mtDNA Assays

Mitochondrial and nuclear DNA in cell-free plasma were quantified using previously described methods (24) with a few modifications. Briefly, plasma samples were thawed from storage at −80°C and centrifuged (5,000 x g, 10 minutes, 4 °C; Eppendorf 5427R with rotor FA-45-48-11; Eppendorf, Enfield, CT). Supernatants were transferred to 96-well plates and stored at −80°C until analysis. After thawing plates, samples were thermolyzed overnight on replicate 96-well plates. Replicate lysates were analyzed in triplicates on 384-well plates using TaqMan chemistry-based real time quantitative polymerase chain reactions (qPCR) targeting mitochondrial gene ND1 and nuclear gene B2M. The medians of triplicate cycle threshold (C_T_) values of samples were compared to those of serial dilutions of DNA standards to determine absolute copy numbers of target genes. Average PCR efficiencies for ND1 and B2M were 96.1% and 94.5%, respectively. The average coefficients of variation of natural log transformed ND1 and B2M copy number between replicates were 2.6% and 6.8%. Copy numbers were adjusted by plate-specific correction factors calculated from measurements of reference standards to correct for batch effects. Detailed information about these methods is available in the supplemental information.

### 6. Statistical analysis

Statistical analyses were conducted using GraphPad Prism (version 9.4.1) and R Software (version 4.2.2 and 4.3.0). The change in GDF15 between pairs of visits was calculated by subtracting the GDF15 levels measured during the former visit from those measured during the later visit. Non-parametric signed-rank Wilcoxon paired t-test was used to compare levels between visits. In EPI, due to challenges in recruiting participants during early pregnancy, difficulties with blood collection, and postpartum dropouts, no participants had data available for all four time points. Considering the non-normal distribution of cf-mtDNA and GDF15 levels and the imbalance in sample sizes, non-parametric pairwise comparisons were utilized to make the most of the available data. This method ensures clear and meaningful insights into changes between specific time points while reducing the impact of missing data and small sample sizes on the reliability and validity of the findings. Spearman rank correlations were used to assess continuous associations. Non-parametric Mann-Whitney t-test was used to assess group difference.

## Availability of data and material

GDF15 immunohistochemistary data of multiple tissue for supplemental figure 1 can be retrieved from Human Protein Atlas https://www.proteinatlas.org/ENSG00000130513-GDF15/tissue. All other data generated in this project will be made available to researchers upon request.

## Author’s Contribution

C.T. and E.A.D. conceived and supervised this research project. C.M. designed the EPI study and supervised data collection. E.A.D. designed the BABIP study and supervised data collection. Q.H., S.A., and D.S. performed the GDF15 and cf-mtDNA assays. Q.H. performed statistical analyses and prepared the figures. C.T. and Q.H. drafted the manuscript. E.A.D., M.P. and F.H. advised on manuscript and figure preparation. All authors reviewed, commented and edited the final version of the manuscript.

