## Supplemental Information for "Associations between prenatal distress, mitochondrial health, and gestational age: findings from two pregnancy studies in the USA and Turkey"

**Supplemental methods:**

Mitochondrial and nuclear DNA in cell-free plasma were quantified using the method described in Michelson et al. 2023^1^ with the following changes and additional details. In instances where the cited method indicates that plates are to be vortexed, they were instead shaken at 1,500 rpm for 1 minute on an OrbiShaker MP (Benchmark Scientific, Sayreville, NJ) and then briefly centrifuged using an Axygen plate centrifuge (Corning Life Sciences, Corning, NY). Models of thermal cyclers used for lysis were either T100 (Bio-Rad, Hercules, CA) or ProFlex PCR System (Applied Biosystems, Waltham, MA). Absolute copy numbers of target genes in samples from the EPI study were determined by comparing their $C_{T}$ measurements versus those measured in serial dilutions of DNA extracted from human fibroblasts (hFB), as described in the cited method, while samples from the BABIP study were quantified using a chimeric ND1-B2M synthetic DNA fragment (sND1-B2M; gBlocks® gene fragment, Integrated DNA Technologies, Coralville, IA), also described in the cited method. The absolute copy numbers of target genes in sND1-B2M were determined by quantifying it using the hFB standard, which was absolutely quantified by digital droplet polymerase chain reaction (Figure 1). Instead of quantifying based on the averages of triplicate $C_{T}$ measurements after discarding outliers, as described in the cited method, the medians of triplicate $C_{T}$ measurements ($C_{Tmed})$ without outlier removal were used. For each qPCR plate, linear regressions were fit to $C_{Tmed}$ versus natural log-transformed (LN) copy number in the serially diluted standard for each target gene. Absolute copy numbers of target genes in samples were calculated using the following formula:

$$Copy per reaction=e^{\frac{C_{Tmed}-b}{m}}$$

Where $b$ and $m$ are the slope and intercept of the linear regression of the standard curve for the corresponding qPCR plate and target gene. A target gene was considered below the limit of detection in a reaction when its copy number was three or fewer^2^. Copy per reaction values were transformed to copy per µL saliva by correcting for dilution of samples into lysis and qPCR buffers. Eight reference standards, created by pooling samples from the Picard lab biobank, were loaded on each 96-well plate alongside saliva samples prior to lysis to assess and correct for plate effects. The measurements of these standards were used to calculate correction factors that were specific to each pair of replicate plates as follows:

$${cf}_{p}=\frac{\bar{LN(ND1)}_{p}}{\bar{LN(ND1)}_{q}}$$

Where ${cf}_{p}$ is the correction factor for plate $p$, and $\bar{LN(ND1)}_{p}$ and $\bar{LN(ND1)}_{q}$ are the averages of LN ND1 copy number per µL in reference standards on plate $p$ and on 56 previously run plates, respectively (Figures 2). All measurements of ND1 and B2M in samples were then adjusted by dividing their LN copy number values by the correction factor for their corresponding plate. Adjusted values were used in subsequent statistical analyses. On one pair of replicate plates loaded with BABIP plasma samples, the median C.V. between replicate measurements (5.9%) exceeded our threshold of 4.0% for repeating the analysis. The median C.V. for this set of samples was reduced to 5.7% upon repeat analysis. Values from the repeat measurements were used in subsequent analyses.

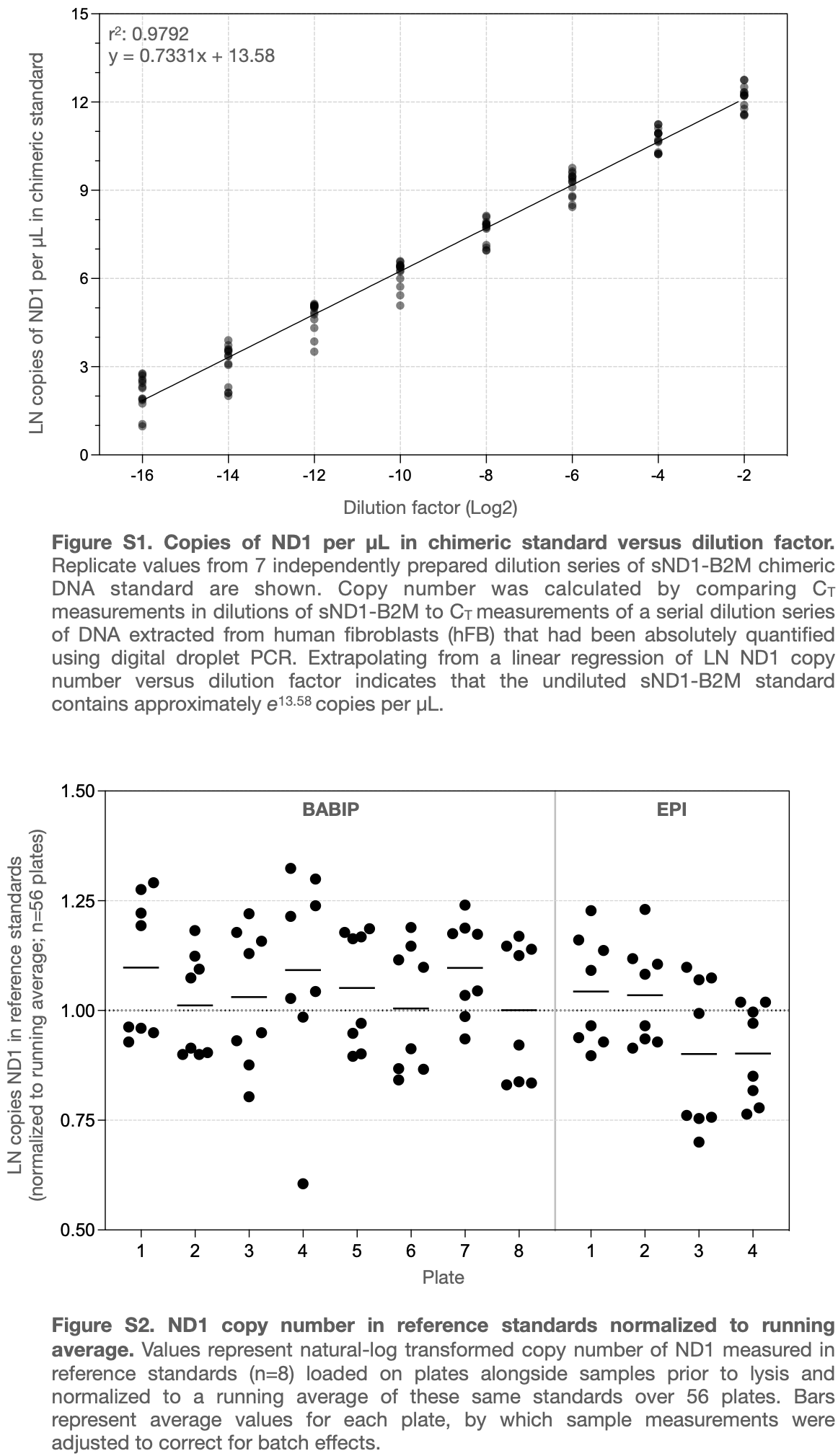

1.	Michelson, J., Rausser, S., Peng, A., Yu, T., Sturm, G., Trumpff, C., Kaufman, B.A., Rai, A.J., and Picard, M. (2023). MitoQuicLy: A high-throughput method for quantifying cell-free DNA from human plasma, serum, and saliva. Mitochondrion *71*, 26-39. 10.1016/j.mito.2023.05.001.

2.	Bustin, S.A., Benes, V., Garson, J.A., Hellemans, J., Huggett, J., Kubista, M., Mueller, R., Nolan, T., Pfaffl, M.W., Shipley, G.L., et al. (2009). The MIQE guidelines: minimum information for publication of quantitative real-time PCR experiments. Clin Chem *55*, 611-622. 10.1373/clinchem.2008.112797.
