## Supplemental Tables for "Associations between prenatal distress, mitochondrial health, and gestational age: findings from two pregnancy studies in the USA and Turkey"

**Supplemental Table 1: Associations between maternal plasma cf-mtDNA levels and maternal characteristics, birth outcomes and prenatal distress.**

|  | | Early 2nd Trimester (EPI-only) | | | Late 2nd Trimester  (EPI & BABIP) | | | 3rd Trimester  (EPI & BABIP) | | | Postpartum  (EPI-only) | | |
| --- | --- | --- | --- | --- | --- | --- | --- | --- | --- | --- | --- | --- | --- |
|  | | 13-18 weeks | | | 19-30 weeks | | | 27-39 weeks | | | 16-56 weeks | | |
|  | | r | p | n | r | p | n | r | p | n | r | p | n |
| **cf-mtDNA** | | | | | | | | | | | | | |
| **Maternal characteristics** | | | | | | | | | | | | | |
| **Age** | | | | | | | | | | | | | |
| EPI  BABIP | | -0.20 | 0.38 | 21 | -0.050 | 0.56 | 156 | -0.08 | 0.35 | 141 | -0.01 | 0.97 | 28 |
|  |  |  |  |  | -0.10 | 0.19 | 183 | 0.05 | 0.60 | 119 |  |  |  |
| **Pre-pregnancy BMI** | | | | | | | | | | | | | |
| EPI  BABIP | | -0.094 | 0.69 | 21 | 0.018 | 0.83 | 155 | 0.00047 | 0.99 | 141 | -0.24 | 0.22 | 28 |
|  |  |  |  |  | -0.061 | 0.41 | 180 | -0.060 | 0.52 | 117 |  |  |  |
| **Pregnancy outcomes** | | | | | | | | | | | | | |
| **Gestational age at birth** | | | | | | | | | | | | | |
| EPI  BABIP | | 0.035 | 0.88 | 20 | 0.046 | 0.57 | 153 | 0.11 | 0.20 | 140 | 0.27 | 0.16 | 28 |
|  |  |  |  |  | 0.0079 | 0.93 | 135 | -0.09 | 0.38 | 99 |  |  |  |
| **Prenatal distress** | | | | | | | | | | | | | |
| **Depression** | | | | | | | | | | | | | |
| EPI (HAM-D) | | -0.56 | **0.032** | 15 | -0.00041 | 0.96 | 154 | 0.0017 | 0.98 | 141 | 0.20 | 0.39 | 20 |
| BABIP (BDI-II) | |  |  |  | 0.021 | 0.78 | 180 | 0.17 | 0.077 | 110 |  |  |  |
| BABIP (CESD) | |  |  |  | -0.054 | 0.47 | 181 | 0.064 | 0.51 | 110 |  |  |  |
| **Anxiety** | | | | | | | | | | | | | |
| EPI (HAM-A) | | -0.51 | **0.055** | 15 | -0.030 | 0.71 | 154 | -0.0076 | 0.93 | 136 | 0.38 | 0.10 | 20 |
| BABIP (STAI-S) | |  |  |  | 0.12 | 0.087 | 179 | -0.021 | 0.83 | 108 |  |  |  |
| **Perceived Stress (PSS)** | | | | | | | | | | | | | |
| EPI | | -0.72 | **0.0031** | 15 | -0.052 | 0.52 | 153 | 0.14 | 0.11 | 136 | -0.13 | 0.57 | 23 |
| BABIP | |  |  |  | 0.029 | 0.70 | 181 | 0.071 | 0.46 | 109 |  |  |  |

P-values and effect sizes from Spearman’s rank correlation. BMI: Body mass index; HAM-D: Hamilton Depression Rating Scale; BDI-II: Beck’s Depression Inventory-II; CESD: Center for Epidemiological Studies Depression; HAM-A: Hamilton Anxiety Rating Scale; STAI-S: State-Trait Anxiety Inventory-State; PSS: Perceived Stress Scale. This table included all participants.

**Supplemental Table 2: Associations between maternal plasma GDF15 levels and maternal characteristics, birth outcomes and prenatal distress.**

|  | | Early 2nd Trimester (EPI-only) | | | Late 2nd Trimester  (EPI & BABIP) | | | 3rd Trimester  (EPI & BABIP) | | | Postpartum  (EPI-only) | | |
| --- | --- | --- | --- | --- | --- | --- | --- | --- | --- | --- | --- | --- | --- |
|  | | 13-18 weeks | | | 19-30 weeks | | | 27-39 weeks | | | 16-56 weeks | | |
|  | | r | p | n | r | p | n | r | p | n | r | p | n |
| **GDF15** | | | | | | | | | | | | | |
| **Maternal characteristics** | | | | | | | | | | | | | |
| **Age** | | | | | | | | | | | | | |
| EPI | | 0.062 | 0.7900 | 21 | 0.00044 | 0.99 | 156 | -0.12 | 0.16 | 141 | 0.47 | **0.016** | 26 |
| BABIP | |  |  |  | -0.14 | 0.054 | 183 | -0.18 | **0.049** | 119 |  |  |  |
| **Pre-pregnancy BMI** | | | | | | | | | | | | | |
| EPI | | 0.03 | 0.9043 | 21 | -0.17 | **0.039** | 155 | -0.15 | 0.068 | 141 | 0.14 | 0.50 | 26 |
| BABIP | |  |  |  | -0.13 | 0.090 | 180 | -0.05 | 0.57 | 117 |  |  |  |
| **Birth outcomes** | | | | | | | | | | | | | |
| **Gestational age at birth** | | | | | | | | | | | | | |
| EPI | | 0.17 | 0.49 | 20 | -0.22 | **0.0075** | 153 | -0.056 | 0.51 | 140 | 0.31 | 0.13 | 26 |
| BABIP | |  |  |  | 0.05 | 0.60 | 135 | 0.0082 | 0.94 | 99 |  |  |  |
| **Prenatal distress** | | | | | | | | | | | | | |
| **Depression** | | | | | | | | | | | | | |
| EPI (HAM-D) | | -0.44 | 0.086 | 16 | 0.061 | 0.45 | 154 | -0.16 | 0.063 | 134 | 0.029 | 0.91 | 19 |
| BABIP (BDI-II) | |  |  |  | 0.074 | 0.36 | 153 | -0.12 | 0.37 | 62 |  |  |  |
| BABIP (CESD) | |  |  |  | -0.012 | 0.88 | 152 | -0.081 | 0.55 | 57 |  |  |  |
| **Anxiety** | | | | | | | | | | | | | |
| EPI (HAM-A) | | -0.50 | 0.0526 | 16 | -0.059 | 0.47 | 154 | -0.16 | 0.055 | 136 | -0.19 | 0.44 | 19 |
| BABIP (STAI-S) | |  |  |  | 0.12 | 0.17 | 140 | 0.22 | 0.099 | 56 |  |  |  |
| **Perceived Stress (PSS)** | | | | | | | | | | | | | |
| EPI | | -0.35 | 0.1827 | 16 | 0.039 | 0.63 | 153 | 0.036 | 0.68 | 136 | -0.36 | 0.11 | 21 |
| BABIP | |  |  |  | -0.044 | 0.59 | 152 | 0.0052 | 0.96 | 90 |  |  |  |

P-values and effect sizes from Spearman’s rank correlation. BMI: Body mass index; HAM-D: Hamilton Depression Rating Scale; BDI-II: Beck’s Depression Inventory; CESD: Center for Epidemiological Studies Depression; HAM-A: Hamilton Anxiety Rating Scale; STAI-S: State-Trait Anxiety Inventory-State; PSS: Perceived Stress Scale. This table included all participants.

**Supplemental Table 3: Differences in maternal plasma cf-mtDNA and GDF15 levels by neonatal sex.**

|  | | Early 2nd Trimester  (EPI-only) | | | | Late 2nd Trimester  (EPI & BABIP) | | | | 3rd Trimester  (EPI & BABIP) | | | | Postpartum  (EPI-only) | | | |
| --- | --- | --- | --- | --- | --- | --- | --- | --- | --- | --- | --- | --- | --- | --- | --- | --- | --- |
|  | | 13-18 weeks | | | | 19-30 weeks | | | | 27-39 weeks | | | | 16-56 weeks | | | |
|  | | Conc. | SD | n | p | Conc. | SD | n | p | Conc. | SD | n | p | Conc. | SD | n | p |
| **Neonatal sex** | | | | | | | | | | | | | | | | | |
| **cf-mtDNA** | | | | | | | | | | | | | | | | | |
| EPI | F | 9629 | 22721 | 15 | 0.61 | 3434 | 5257 | 74 | 0.40 | 3919 | 5468 | 73 | 0.38 | 8916 | 11163 | 12 | 0.38 |
|  | M | 2379 | 2232 | 5 |  | 4429 | 7169 | 79 |  | 3245 | 4670 | 67 |  | 4185 | 6797 | 14 |  |
| BABIP | F |  |  |  | | 2276 | 2350 | 68 | 0.68 | 2551 | 2325 | 50 | 0.38 |  |  |  | |
|  | M |  |  |  | | 2553 | 2776 | 85 |  | 2344 | 2292 | 59 |  |  |  |  | |
| **GDF15** | | | | | | | | | | | | | | | | | |
| EPI | F | 14320 | 8712 | 15 | 0.67 | 17271 | 11661 | 74 | 0.42 | 36772 | 31879 | 73 | 0.59 | 446 | 173.9 | 12 | 0.63 |
|  | M | 17182 | 12156 | 5 |  | 16327 | 12109 | 79 |  | 32755 | 24531 | 67 |  | 681.3 | 1149 | 14 |  |
| BABIP | F |  |  |  | | 14929 | 8085 | 68 | 0.11 | 27746 | 15553 | 50 | 0.082 |  |  |  | |
|  | M |  |  |  | | 12568 | 5353 | 85 |  | 23288 | 14924 | 59 |  |  |  |  | |

P-values and effect sizes from non-parametric Mann-Whitney t-test. Conc.: Concentration (cf-mtDNA measured in copies/uL, GDF15 measured in pg/mL); SD: standard deviation; F: Female; M: Male. This table included all participants.

**Supplemental Table 4: Differences in maternal plasma cf-mtDNA and GDF15 levels by perinatal complications.**

|  | | Late 2nd Trimester  (EPI & BABIP) | | | | 3rd Trimester  (EPI & BABIP) | | | |
| --- | --- | --- | --- | --- | --- | --- | --- | --- | --- |
|  | | 19-30 weeks | | | | 27-39 weeks | | | |
|  | | Conc. | SD | n | p | Conc. | SD | n | p |
| **cf-mtDNA** | | | | | | | | |  |
| **Perinatal complications** | | | | | | | | |  |
| **Preeclampsia** | | | | | | | | |  |
| EPI | No | 4182 | 6738 | 150 | 0.95 | 3673 | 5186 | 136 | 0.51 |
|  | Yes | 2123 | 1240 | 6 |  | 2669 | 1238 | 5 |  |
| BABIP | No | 2339 | 2443 | 187 | NA | 2193 | 1864 | 118 | NA |
|  | Yes | 955.0 | 214.2 | 2 |  | 11246 | 4525 | 2 |  |
| **Preterm birth** | | | | | | | | | |
| EPI | No | 3863 | 6472 | 146 | **0.044** | 3687 | 5144 | 138 | 0.43 |
|  | Yes | 7600 | 8123 | 10 |  | 1372 | 551.6 | 3 |  |
| BABIP | No | 2359 | 2471 | 181 | 0.33 | 2340 | 2239 | 115 | 0.90 |
|  | Yes | 1541 | 1197 | 8 |  | 2443 | 2035 | 5 |  |
| **Gestational diabetes** | | | | | | | | | |
| EPI | No | 4048 | 6694 | 151 | **0.022** | 3612 | 5121 | 138 | 0.30 |
|  | Yes | 5771 | 3909 | 5 |  | 4825 | 4688 | 3 |  |
| BABIP | No | 2514 | 2678 | 104 | 0.092 | 2396 | 2470 | 76 | **0.013** |
|  | Yes | 2196 | 1678 | 15 |  | 3420 | 2024 | 12 |  |
| **GDF15** | | | | | | | | |  |
| **Perinatal complications** | | | | | | | | |  |
| **Preeclampsia** | | | | | | | | |  |
| EPI | No | 17144 | 11926 | 150 | **0.017** | 34237 | 26386 | 136 | 0.35 |
|  | Yes | 8348 | 4596 | 6 |  | 47842 | 68741 | 5 |  |
| BABIP | No | 13678 | 6763 | 187 | NA | 25671 | 15108 | 118 | NA |
|  | Yes | 6970 | 3451 | 2 |  | 13557 | 5360 | 2 |  |
| **Preterm birth** | | | | | | | | | |
| EPI | No | 16418 | 11732 | 146 | 0.069 | 34899 | 28653 | 138 | 0.89 |
|  | Yes | 22472 | 12664 | 10 |  | 26428 | 23493 | 3 |  |
| BABIP | No | 13589 | 6768 | 181 | 0.88 | 25672 | 15328 | 115 | 0.68 |
|  | Yes | 14012 | 7185 | 8 |  | 20809 | 5628 | 5 |  |
| **Gestational diabetes** | | | | | | | | | |
| EPI | No | 17003 | 11965 | 151 | 0.28 | 35003 | 28730 | 138 | 0.42 |
|  | Yes | 10840 | 4671 | 5 |  | 21647 | 9439 | 3 |  |
| BABIP | No | 13518 | 6897 | 104 | 0.34 | 24308 | 14108 | 76 | 0.78 |
|  | Yes | 11897 | 5676 | 15 |  | 28688 | 18800 | 12 |  |

P-values and effect sizes from non-parametric Mann-Whitney t-test. Preterm birth was defined as delivery before 37 completed weeks of gestation; and gestational diabetes was diagnosed based on standard oral glucose tolerance test criteria during pregnancy. Conc.: Concentration (cf-mtDNA measured in copies/uL, GDF15 measured in pg/mL); SD: standard deviation; NA: sample size too small for statistically analysis. This table included all participants.

**Supplemental Table 5: Descriptive statistics of maternal plasma cf-mtDNA and GDF15 levels by perinatal distress measure categories**

|  | | Early 2nd Trimester (EPI-only) | | | Late 2nd Trimester  (EPI & BABIP) | | | 3rd Trimester  (EPI & BABIP) | | | Postpartum  (EPI-only) | | |
| --- | --- | --- | --- | --- | --- | --- | --- | --- | --- | --- | --- | --- | --- |
|  | | 13-18 weeks | | | 19-30 weeks | | | 27-39 weeks | | | 16-56 weeks | | |
|  | | Conc. | SD | n | Conc. | SD | n | Conc. | SD | n | Conc. | SD | n |
| **Prenatal Distress** | | | | | | | | | | | | | |
| **Depression** | | | | | | | | | | | | | |
| **EPI (HAM-D)** | | | | | | | | | | | | | |
| cf-mtDNA | Mild-none | 9727 | 24114 | 13 | 4486 | 7119 | 129 | 3832 | 5409 | 121 | 1157 | 9472 | 18 |
|  | Moderate |  |  |  | 2264 | 2124 | 15 | 4186 | 2625 | 7 | 24613 |  | 1 |
|  | Severe | 1084 | 983.3 | 3 | 2585 | 3832 | 10 | 2625 | 1157 | 6 | 307.0 |  | 1 |
| GDF15 | Mild-none | 14675 | 9304 | 13 | 16481 | 11214 | 129 | 36261 | 30253 | 120 | 696.1 | 1032 | 17 |
|  | Moderate |  |  |  | 14424 | 6586 | 15 | 22936 | 10785 | 7 | 302.8 |  | 1 |
|  | Severe | 10918 | 8113 | 3 | 15402 | 7791 | 10 | 25362 | 8536 | 7 | 410.6 |  | 1 |
| **BABIP (BDI-II)** | | | | | | | | | | | | | |
| cf-mtDNA | Mild-none |  |  |  | 2423 | 2540 | 169 | 2207 | 1855 | 105 |  |  |  |
|  | Moderate |  |  |  | 1453 | 940.2 | 7 | 5044 | 5375 | 5 |  |  |  |
|  | Severe |  |  |  | 1870 | 1180 | 4 |  |  |  |  |  |  |
| GDF15 | Mild-none |  |  |  | 13801 | 6850 | 169 | 25781 | 15713 | 105 |  |  |  |
|  | Moderate |  |  |  | 14971 | 7248 | 7 | 17805 | 3945 | 5 |  |  |  |
|  | Severe |  |  |  | 10650 | 5460 | 4 |  |  |  |  |  |  |
| **BABIP (CESD)** | | | | | | | | | | | | | |
| cf-mtDNA | <16 |  |  |  | 2511 | 2720 | 133 | 2277 | 1961 | 81 |  |  |  |
|  | >=16 |  |  |  | 1921 | 1569 | 48 | 2500 | 2695 | 29 |  |  |  |
| GDF15 | <16 |  |  |  | 13751 | 6856 | 133 | 24382 | 12583 | 81 |  |  |  |
|  | >=16 |  |  |  | 13551 | 6918 | 48 | 28897 | 21578 | 29 |  |  |  |
| **BABIP(Self-reported psychiatric disorder diagnosis)** | | | | | | | | | | | | | |
| cf-mtDNA | No |  |  |  | 2308 | 2403 | 183 | 2398 | 2255 | 115 |  |  |  |
|  | Yes |  |  |  | 2847 | 3514 | 6 | 1106 | 254.7 | 5 |  |  |  |
| GDF15 | No |  |  |  | 13627 | 6808 | 183 | 25539 | 15123 | 115 |  |  |  |
|  | Yes |  |  |  | 13001 | 5840 | 6 | 23862 | 15309 | 5 |  |  |  |
| **Anxiety** | | | | | | | | | | | | | |
| **EPI (HAM-A)** | | | | | | | | | | | | | |
| cf-mtDNA | Mild-none | 11221 | 26112 | 11 | 4879 | 7761 | 91 | 3373 | 5203 | 84 | 3969 | 9348 | 15 |
|  | Moderate | 1744 | 545.5 | 3 | 3194 | 4757 | 48 | 3992 | 5423 | 44 | 10608 | 9785 | 3 |
|  | Severe | 518.3 | 118.9 | 2 | 2752 | 3470 | 15 | 3779 | 2891 | 8 | 12460 | 17187 | 2 |
| GDF15 | Mild-none | 14660 | 9601 | 11 | 16605 | 11284 | 91 | 39224 | 32465 | 84 | 762.4 | 1132 | 14 |
|  | Moderate | 16502 | 8220 | 3 | 16056 | 10485 | 48 | 27928 | 21780 | 43 | 386.7 | 145.8 | 3 |
|  | Severe | 6383 | 2877 | 2 | 14317 | 6653 | 15 | 27757 | 9060 | 9 | 356.7 | 76.23 | 2 |
| **BABIP (STAI-S)** | | | | | | | | | | | | | |
| cf-mtDNA | Mild-non |  |  |  |  |  |  |  |  |  |  |  |  |
|  | Moderate |  |  |  | 2446 | 2321 | 37 | 2468 | 2653 | 26 |  |  |  |
|  | Severe |  |  |  | 2339 | 2540 | 142 | 2277 | 2039 | 82 |  |  |  |
| GDF15 | Mild-non |  |  |  |  |  |  |  |  |  |  |  |  |
|  | Moderate |  |  |  | 13739 | 6795 | 37 | 27157 | 12117 | 26 |  |  |  |
|  | Severe |  |  |  | 13739 | 6927 | 142 | 24744 | 16074 | 82 |  |  |  |
| **BABIP (Self-reported psychiatric disorder diagnosis)** | | | | | | | | | | | | | |
| cf-mtDNA | No |  |  |  | 2336 | 2504 | 169 | 2294 | 2012 | 107 |  |  |  |
|  | Yes |  |  |  | 2231 | 1787 | 20 | 2757 | 3614 | 13 |  |  |  |
| GDF15 | No |  |  |  | 13568 | 6812 | 169 | 25415 | 14825 | 107 |  |  |  |
|  | Yes |  |  |  | 13939 | 6530 | 20 | 25919 | 17619 | 13 |  |  |  |

P-values and effect sizes from Non-parametric Mann-Whitney t-test. HAM-D: Hamilton Depression Rating Scale, 0-13 mild-none, 14-17 moderate, >17 severe; BDI-II: Beck’s Depression Inventory-II, 0-19 mild-non, 20-28 moderate, 29-63 severe; CESD: Center for Epidemiological Studies Depression; HAM-A: Hamilton Anxiety Rating Scale, 0-14 mild-none, 15-24 moderate, >24 severe; STAI-S: State-Trait Anxiety Inventory-State, 20-37 mild-non, 38-44 moderate, 45-80 severe. This table included all participants.

**Supplemental Table 6: Associations between change in maternal plasma GDF15 levels and maternal characteristics, birth outcomes and prenatal distress across respective time points.**

|  | | GDF15 Change from  Early to Late 2^nd^ Trimester  (EPI-only) | | | GDF15 Change from  Early 2^nd^ to 3^rd^ Trimester  (EPI-only) | | |
| --- | --- | --- | --- | --- | --- | --- | --- |
|  | | r | p | n | r | p | n |
| **Maternal characteristics** | | | | | | | |
| **Age** | | | | | | | |
| EPI | | 0.47 | 0.18 | 10 | 0.16 | 0.54 | 17 |
| **Pre-pregnancy BMI** | | | | | | | |
| EPI | | 0.21 | 0.55 | 10 | 0.055 | 0.83 | 17 |
| **Birth outcomes** | | | | | | | |
| **Gestational age at birth** | | | | | | | |
| EPI | | 0.43 | 0.21 | 10 | 0.17 | 0.52 | 17 |
| **Prenatal distress measured at early pregnancy (EPI: Visit 1)** | | | | | | | |
| **Depression** | | | | | | | |
| EPI (HAM-D) | | 0.19 | 0.59 | 10 | 0.52 | 0.085 | 12 |
| **Anxiety** | | | | | | | |
| EPI (HAM-A) | | 0.042 | 0.91 | 10 | 0.54 | 0.074 | 12 |
| **Perceived Stress** | | | | | | | |
| EPI | | 0.14 | 0.71 | 10 | 0.31 | 0.33 | 12 |
|  | | GDF15 Change from  Late 2^nd^ to 3^rd^ Trimester  (EPI & BABIP) | | | GDF15 Change from  Late 2^nd^ Trimester to Postpartum  (EPI-only) | | |
|  | | r | p | n | r | p | n |
| **Maternal characteristics** | | | | | | | |
| **Age** | | | | | | | |
| EPI | | -0.23 | **0.0089** | 127 | -0.18 | 0.40 | 24 |
| BABIP | | -0.072 | 0.44 | 118 |  |  |  |
| **Pre-pregnancy BMI** | | | | | | | |
| EPI | | -0.036 | 0.69 | 124 | 0.31 | 0.15 | 24 |
| BABIP | | -0.0051 | 0.96 | 117 |  |  |  |
| **Birth outcomes** | | | | | | | |
| **Gestational age at birth** | | | | | | | |
| EPI | | 0.13 | 0.13 | 126 | 0.14 | 0.53 | 24 |
| BABIP | | 0.015 | 0.89 | 98 |  |  |  |
| **Prenatal distress measured at late 2^nd^ trimester** | | | | | | | |
| **Depression** | | | | | | | |
| EPI (HAM-D) | | -0.042 | 0.64 | 126 | 0.33 | 0.13 | 23 |
| BABIP (BDI-II) | | -0.10 | 0.29 | 114 |  |  |  |
| BABIP (CESD) | | -0.16 | 0.095 | 111 |  |  |  |
| **Anxiety** | |  |  |  |  |  |  |
| EPI (HAM-A) | | 0.0038 | 0.97 | 126 | 0.21 | 0.34 | 23 |
| BABIP (STAI-S) | | -0.012 | 0.90 | 112 |  |  |  |
| **Perceived Stress (PSS)** | |  |  |  |  |  |  |
| EPI | | 0.040 | 0.66 | 126 | 0.27 | 0.21 | 23 |
| BABIP | | -0.010 | 0.91 | 117 |  |  |  |
|  | | GDF15 Change from  3^rd^ Trimester to Postpartum  (EP only) | | |  | | |
|  | | r | p | n |  |  |  |
| **Maternal characteristics** | | | | | | | |
| **Age** | |  |  |  |  |  |  |
| EPI | | -0.032 | 0.88 | 24 |  |  |  |
| **Pre-pregnancy BMI** | |  |  |  |  |  |  |
| EPI | | 0.41 | **0.044** | 24 |  |  |  |
| **Birth outcomes** | |  |  |  |  |  |  |
| **Gestational age at birth** | | -0.075 | 0.72 | 24 |  |  |  |
| EPI | |  |  |  |  |  |  |
| **Prenatal distress measured at 3^rd^ trimester** | | | | | | | |
| **Depression** | | | | | | | |
| EPI (HAM-D) | | 0.59 | **0.0040** | 22 |  |  |  |
| **Anxiety** | | | | | | | |
| EPI (HAM-A) | | 0.26 | 0.24 | 23 |  |  |  |
| **Perceived Stress** | | | | | | | |
| EPI | | 0.43 | **0.042** | 23 |  |  |  |

P-values and effect sizes from Spearman’s rank correlation. BMI: Body mass index; HAM-D: Hamilton Depression Rating Scale; BDI-II: Beck’s Depression Inventory-II; CESD: Center for Epidemiological Studies Depression; HAM-A: Hamilton Anxiety Rating Scale; STAI-S: State-Trait Anxiety Inventory-State; PSS: Perceived Stress Scale. This table included all participants.
