## Supplemental Figures for "Associations between prenatal distress, mitochondrial health, and gestational age: findings from two pregnancy studies in the USA and Turkey"

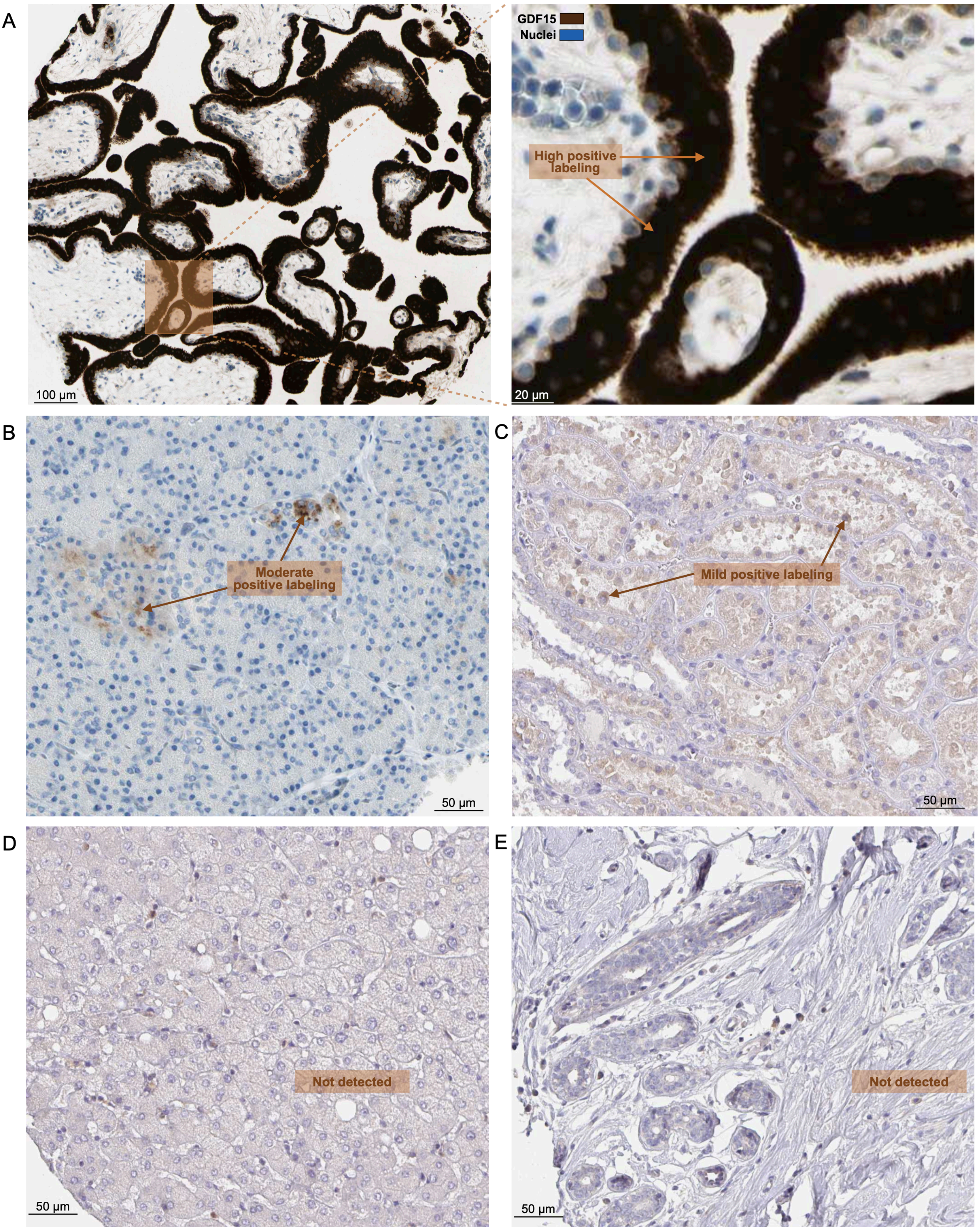


**Figure S1. GDF15 protein immunohistochemistry in different tissues**. Human Protein Atlas (HPA) immunohistochemistry of GDF15 protein in the (**A**) placenta, (**B**) pancreas, (**C**) kidney, and (**D**) liver, (**E**) breast. Subject IDs are #2169 (19-year-old, female) for A, #2032 (35-year-old, female) for B, #2530 (41-year-old, female) for C, #3402 (54-year-old, female) for D, #3856 (27-year-old, female) for E based on sample availability.

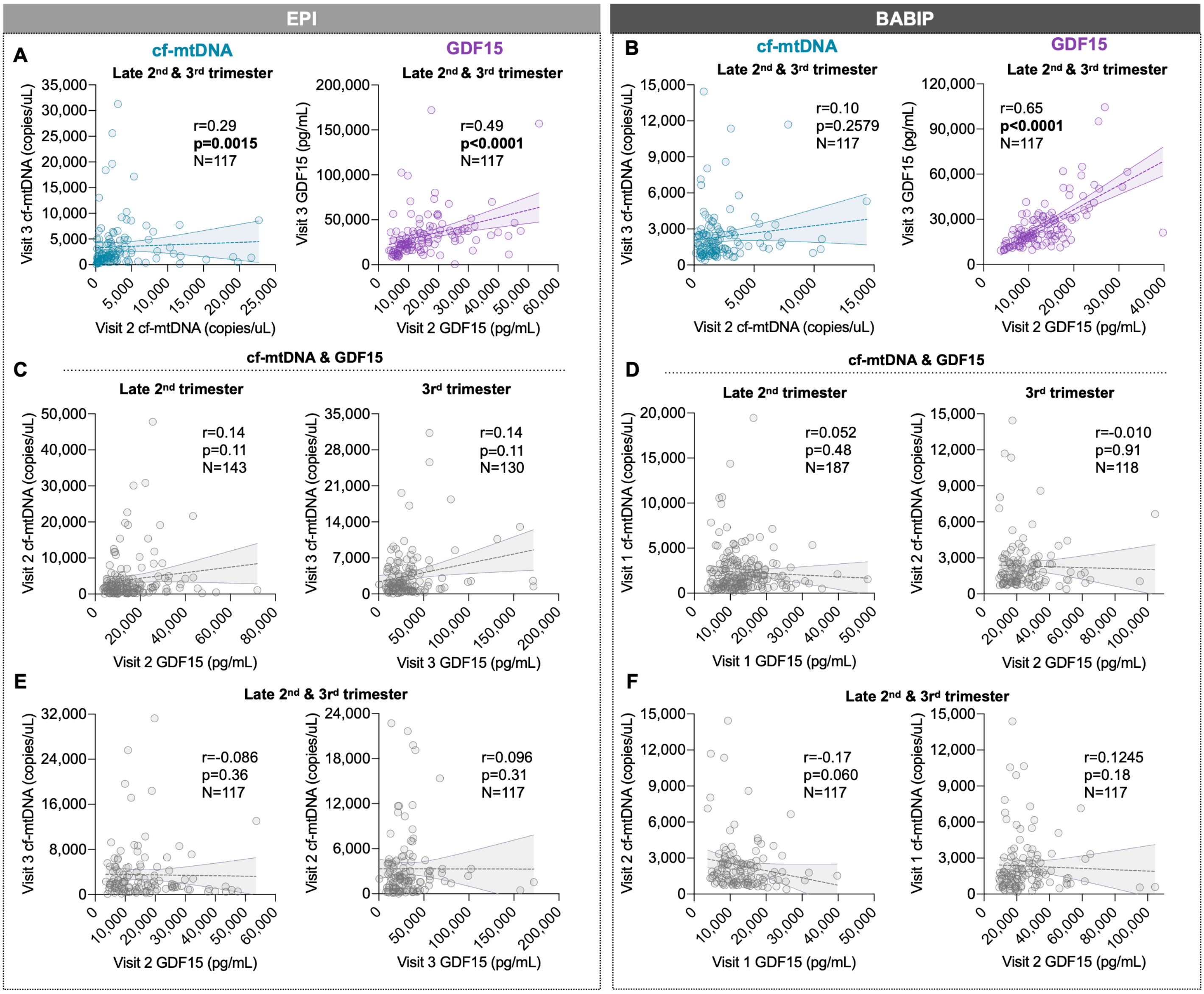


**Figure S2. Association between maternal cf-mtDNA and GDF15 plasma levels across study timepoints.** Association between levels measured at different visits for cf-mtDNA (left) and GDF15 (right) from (**A**) EPI and (**B**) BABIP. Association between cf-mtDNA and GDF15 levels measured in the same visits from (**C**) EPI and (**D**) BABIP. Association between GDF15 and cf-mtDNA levels measured at different visits from (**E**) EPI and (**F**) BABIP. P-values and effect sizes from Spearman's rank correlation. *p<0.05, **p<0.01, ***p<0.001, ****p<0.0001.

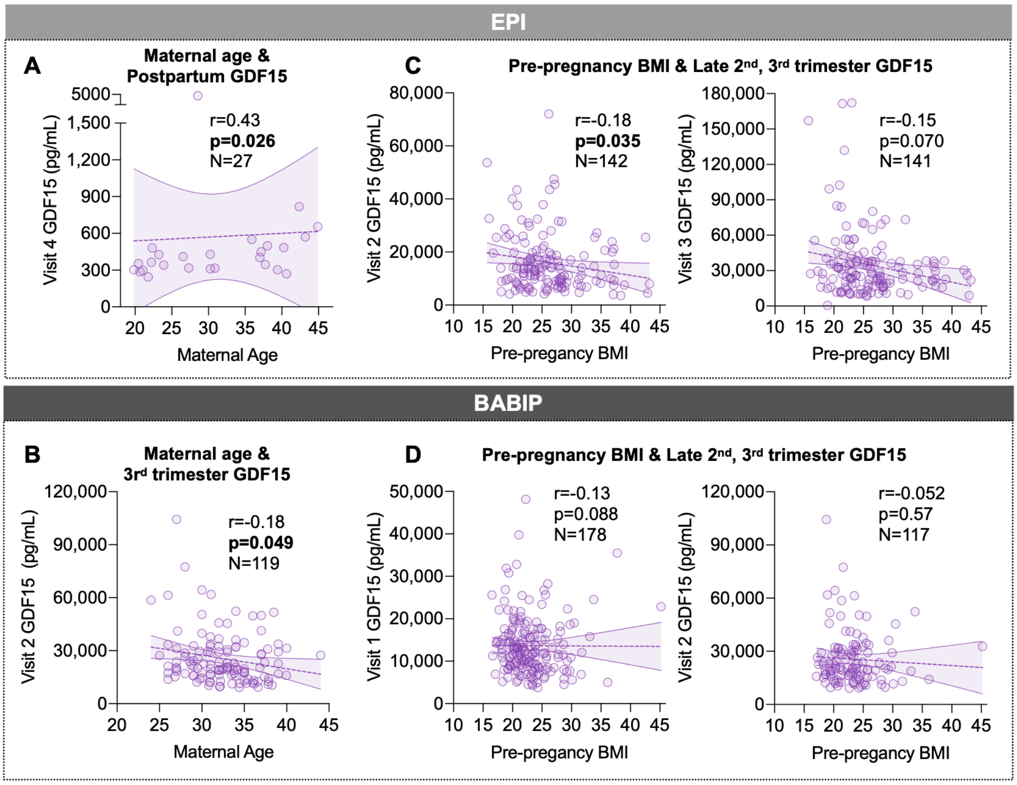


**Figure S3. Associations between GDF15 levels and maternal characteristics.** Associations between maternal age and GDF15 levels at postpartum (**A,** EPI) and late-pregnancy (**B,** BABIP). Associations between pre-pregnancy BMI and GDF15 levels at mid- and late-pregnancy from EPI (**C**) and BABIP (**D**), 13 EPI participants and 2 BABIP participants were removed from the analysis and graphs since they completed their late 2nd trimester visit in their 3rd trimester. P-values and effect sizes from Spearman's rank correlation. *p<0.05, **p<0.01, ***p<0.001, ****p<0.0001.

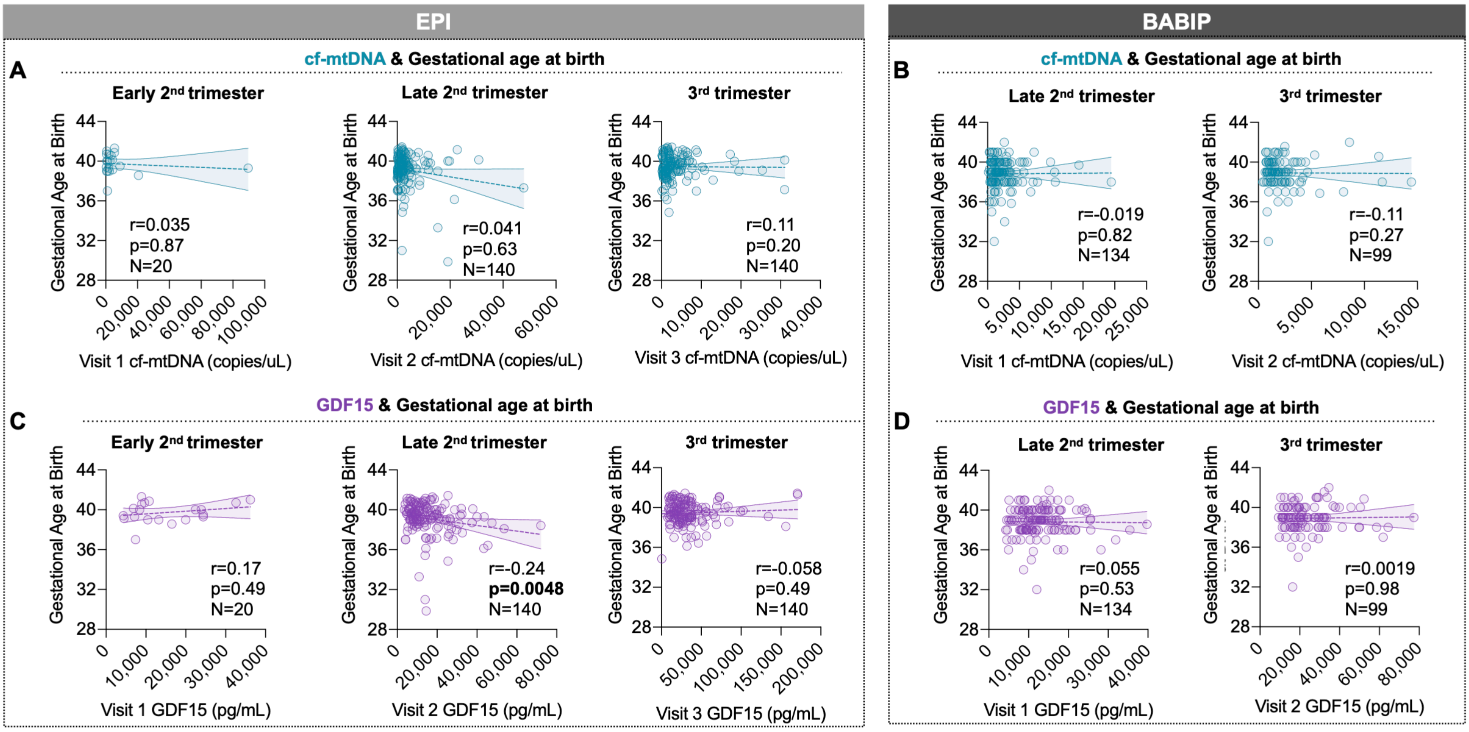


**Figure S4. Association between maternal plasma cf-mtDNA and GDF15 levels and gestational age at birth (raw values).** Associations between gestational age at birth and cf-mtDNA at each visit in (**A**) EPI and (**B**) BABIP. Associations between gestational age at birth and GDF15 at each visit in (**C**) EPI and (**D**) BABIP. 13 EPI participants and 2 BABIP participants were removed from the analysis and graphs since they completed their late 2nd trimester visit in their 3rd trimester. P-values and effect sizes from Spearman's rank correlation. *p<0.05, **p<0.01, ***p<0.001, ****p<0.0001.
